# Auditory experience shapes how listeners exploit perceptual equivalency classes in speech and cope with noise degradation: Evidence from “elliptical speech”

**DOI:** 10.64898/2026.01.02.695202

**Authors:** Gavin M. Bidelman, Zara Eisenhut, Lucy Borowski, Rose Rizzi, David B. Pisoni

## Abstract

We investigated links between perceptual gradiency in phonetic categorization and speech-in-noise (SIN) perception in listeners with varying music backgrounds. Categorization was measured for vowels and stops using phoneme labeling tasks. Speech discrimination and transcription were assessed using “elliptical speech” sentences [Miller and Nicely (1955). *JASA*, 27, 338–352] which use featural substitutions that renders them meaningless in clean conditions but surprisingly improves their recognition under noise. We hypothesized listeners who use broader perceptual equivalency classes at the phoneme and/or sentence level would show better SIN. Perception of elliptical speech was indeed resilient to noise but this elliptical benefit varied with music background; nonmusicians showed larger susceptibility and noise-related benefit from ellipses than musicians. Phoneme categorization and elliptical sentence perception were also associated with QuickSIN performance but in opposite ways depending on musicianship. Broader phonetic category usage was related to better SIN in nonmusicians but poorer SIN in musicians. Findings suggest listeners can use broader/narrower perceptual equivalency classes and top-down inference to cope with noise degradation but this depends on auditory demographics. Musically naïve listeners can use broader phonetic categories to aid SIN perception while expert listeners can use narrower categories in otherwise similar speech contexts.

**PACS numbers:** 43.66, 43.71

## I. INTRODUCTION

Speech contains a larger number of variable acoustic features that impose costs on perception (Mullennix and Pisoni, 1990; Kapadia *et al*., 2023). To reduce perceptual demands and deal with the “invariance problem” in speech, listeners must normalize the signal in various ways (for review, see Johnson, 2005). Categorization is one such process that maps continuous variations in the acoustic input to higher-level abstract phonetic categories to promote perceptual constancy (Harnad, 1987; Goldstone and Hendrickson, 2010). While listeners can exploit category-level information at the phoneme level, they also have simultaneous access to continuous properties of the signal to support its perceptual interpretation (Pisoni and Tash, 1974; Clayards *et al*., 2008; Kapnoula *et al*., 2017; Toscano *et al*., 2018). How they weight these two sources of information influences their so-called “listening strategy” (Kapnoula *et al*., 2017; Rizzi and Bidelman, 2024; Bidelman *et al*., 2025a; Rizzi and Bidelman, 2025). Discrete listeners ignore continuous, surface level features of the speech signal, warping their perception onto a smaller, finite set of category labels. In contrast, gradient listeners retain more within-category signal details resulting in a more linear, one-to-one mapping between signal acoustics and the perceptual space.

Listening strategy can be quantified from categorization paradigms using typical 2AFC phoneme labeling tasks where participants are asked to identify sounds sampled along an acoustic-phonetic continuum (Liberman *et al*., 1957; Pisoni, 1973; Massaro and Cohen, 1983; Bidelman *et al*., 2013; Kong and Edwards, 2016; e.g., Kapnoula *et al*., 2017; e.g., Bidelman *et al*., 2020; Rizzi and Bidelman, 2024). The slope of the resulting identification curve represents the degree of categoricity/gradiency in a listener’s response pattern; shallower slopes reflect more gradient perception, whereas steeper identification reflects a discrete mode of hearing (Sussman, 1993; Hallé *et al*., 2004; Xu *et al*., 2006; Bidelman, 2015).

Emerging evidence suggests that exactly how listeners form categories might benefit another important aspect of receptive communication: speech-in-noise (SIN) recognition. Consequently, the degree to which someone shows more gradient vs. categorical responses in simple phoneme labeling tasks might be closely associated with their SIN performance (Gifford *et al*., 2014; Kapnoula *et al*., 2017; Bidelman *et al*., 2020; Bidelman and Carter, 2023; Myers *et al*., 2024; Rizzi and Bidelman, 2024; Bidelman *et al*., 2025a; Rizzi and Bidelman, 2025). Recent studies have shown more gradient hearing benefits performance in sentence-level SIN recognition (Myers *et al*., 2024; Rizzi and Bidelman, 2024) and “cocktail party” (i.e., speech-on-speech) scenarios (Bidelman *et al*., 2025a). Gradient modes of listening may also facilitate perceptual processing that requires flexibility in category mapping including dialect learning (Wagner *et al*., 2014; McMurray, 2022), recovery from ambiguity (Kapnoula *et al*., 2021b), and coping with indexical variation (McMurray, 2022).

On the contrary, discretizing the signal could make listeners more decisive in their decisions (Tuller *et al*., 1994; Bidelman and Carter, 2023). Under this notion, the natural binning process of categorization might enable category members to “pop out” among a noisy feature space, thereby facilitating speech in noise processing (e.g., Nothdurft, 1991; Pérez-Gay Juárez *et al*., 2019; Bidelman *et al*., 2020). Categories might thus serve as attractor states, which act as a landing point for perception (Eimas and Corbit, 1973; Rozsypal *et al*., 1985). This notion is supported in spoken word recognition, where real words and high-frequency words are more successfully perceived in noise than pseudowords or low-frequency words (e.g., Rosenzweig and Postman, 1957; Pisoni, 1996). Collectively, previous studies highlight an emerging and testable link between phonetic categorization and SIN processing: differences in perceptual gradiency might account for substantial unrecognized variance in SIN perception skills.

Further evidence for categorization-SIN transfer originates from studies examining listeners with auditory expertise and disorders. For example, individuals with high degrees of musicality show sharper identification functions for phoneme labeling (Elmer *et al*., 2012; Bidelman *et al*., 2014; Bidelman and Alain, 2015; Mankel and Bidelman, 2018; Bidelman and Walker, 2019; Mankel *et al*., 2020; Ma *et al*., 2024) and tolerate more severe signal-to-noise ratios (SNR) in degraded speech perception than their musically naïve peers (Parbery-Clark *et al*., 2009b; Zendel and Alain, 2012; Anaya *et al*., 2016; Coffey *et al*., 2017; Mankel and Bidelman, 2018; Yoo and Bidelman, 2019; Bidelman and Yoo, 2020; Benítez-Barrera *et al*., 2022; Hennessy *et al*., 2022; Fang, 2025)^1^. On the contrary, many communication disorders [e.g., specific language impairment (Thibodeau and Sussman, 1979), dyslexia (Werker and Tees, 1987; Noordenbos and Serniclaes, 2015; Gabay *et al*., 2023), auditory processing disorder (Jerger *et al*., 1987)] commonly present with weaker categorization and SIN deficits (Cunningham *et al*., 2001; Warrier *et al*., 2004; Putter-Katz *et al*., 2008). These studies suggest the degree to which an individual is more gradient/continuous in their phoneme perception might be inherently related to their SIN processing abilities.

A limitation of these aforementioned studies is the sole use of isolated phoneme labeling tasks to assess categorization (McMurray *et al*., 2008; Kapnoula *et al*., 2017; Kapnoula *et al*., 2021a; Myers *et al*., 2024; Rizzi and Bidelman, 2024; 2025). While a psychometric slope can quantify how discretely individual phonemes are heard, it provides little insight into how listeners exploit category information at a larger, more ecological level of speech analysis (e.g., sentence-level recognition). Attempts to map isolated phoneme hearing to complex SIN perception (e.g., Kapnoula *et al*., 2017; Kapnoula *et al*., 2021a; Myers *et al*., 2024; Rizzi and Bidelman, 2024) seem dubious given the disparate levels of analysis in these processes. Unique paradigms that allow for the assessment of broad vs. narrow category use during noise-degraded perception of *sentence-level* stimuli would be advantageous to confirm whether category usage and SIN processing are related factors in speech perception (cf. Myers *et al*., 2024; Rizzi and Bidelman, 2024; 2025).

One way to investigate SIN processing in the context of category structure is “elliptical speech” stimuli (Miller and Nicely, 1955). Elliptical speech represents a form of speech distortion where consonants in an equivalence class are swapped with one another, forming an “ellipsis” which degrades the place of articulation information (Miller, 1956). In this featural substitution procedure, stops, fricatives, and nasal consonants for sentence keywords are replaced with new consonants that preserve the same manner and voicing features of the original but transforms place of articulation. As an example, the intact sentence “A *wisp* of *cloud hung* in the *blue air*” under elliptical transformation becomes “A *wist* of *tloud hund* in the *dlue air*” (keywords italicized).

Using elliptical speech, Miller & Nicely (1955) characterized consonant confusions under a wide range of noise masking conditions. They found that place of articulation cues were expectedly confused under signal degradations. Since confusability is a proxy measure of similarity (Miller, 1956), they theorized that based on the pattern of confusion errors, consonants which are more often confused in noise might form perceptual equivalence classes (i.e., broad phonetic categories) that are essentially exchangeable under perceptually degraded conditions. Interestingly, ellipses that were clearly detectable in quiet became surprisingly undetectable amidst noise (i.e., sentences were heard as their “normal”/intact counterpart and the place of articulation swap was not perceived), resulting in improved speech perception performance (Miller and Nicely, 1955; Miller, 1956). Elliptical benefits are generally larger for consonant than vowel sounds (Bond, 1981) and have been reported in subsequent studies in both normal hearing (Quillet *et al*., 1998) and cochlear implant listeners (Herman and Pisoni, 2003). As stated by Miller (1956), “The illusion is heightened if the listener knows what the passage is, for then contextual dependencies fill in the ambiguous distinctions and speech seems clear and distinct (p. 358).” Elliptical speech stimuli are provocative because they suggest that noise can counterintuitively improve speech recognition under states of signal distortion. Presumably, the elliptical effect results from stronger top-down processing (e.g., predictive coding, contextual inference) that must be recruited to cope with signal degradation (Herman and Pisoni, 2003).

Here, we aimed to further examine putative relationships between gradiency in auditory perceptual processing and SIN listening (cf. Kapnoula *et al*., 2017; Bidelman *et al*., 2020; Bidelman and Carter, 2023; Rizzi and Bidelman, 2024; Bidelman *et al*., 2025a; Rizzi and Bidelman, 2025) through the unique lens of elliptical speech stimuli (Quillet *et al*., 1998; Herman and Pisoni, 2003). Our study addressed two primary questions. First, we asked whether isolated phonetic categorization related to SIN processing—does gradient or categorical hearing of tokens confer larger benefit in noise? Second, do listeners who use broader perceptual equivalence classes (at the phoneme and/or sentence levels) perform better in SIN perception? To this end, we measured phoneme categorization (identification labeling) for CV and vowel sounds, SIN recognition (QuickSIN), and elliptical speech perception in clean and noise-degraded conditions. Speech recognition and discrimination were assessed using “elliptical speech” sentences that contained featural substitutions which renders them meaningless under normal (clean) conditions but counterintuitively improves their recognition under noise degradation (Quillet *et al*., 1998; Herman and Pisoni, 2003).

We hypothesized that listeners with more gradient perception at the phoneme level and/or broader category use at the sentence level during elliptical speech would show better SIN perception (e.g., Quillet *et al*., 1998). This would imply more continuous, and arguably more cognitive demanding, modes of perception benefit SIN processing (e.g., Rizzi and Bidelman, 2024; Bidelman *et al*., 2025a). Our sample included listeners with a variety of self-reported music background. This allowed us to assess whether auditory expertise moderates the link between SIN recognition and categorization as implied by prior work (e.g., Coffey *et al*., 2017; Mankel and Bidelman, 2018; Bidelman and Walker, 2019; Hennessy *et al*., 2022). Our findings demonstrate listeners can use broader/narrower perceptual equivalency classes and top-down inference to cope with noise degradation but this link between perceptual gradiency/categoricity in hearing and SIN processing depends on listeners’ music background.

## II. METHOD

### A. Participants

The sample included N=28 young adults (7 male, 21 female; age: *M* = 21.9 ± 5.2 years) recruited from the Indiana University student body and Bloomington community. The sample size was more than 3x larger than previous speech perception studies using identical elliptical speech materials and thus was well powered to detect perceptual differences between intact and elliptical speech (Herman and Pisoni, 2003). All participants exhibited normal hearing sensitivity confirmed via audiometric screening (i.e., < 25 dB HL, octave frequencies 250 -8000 Hz). The majority were strongly right-handed (Oldfield, 1971) (mean laterality index = 59.8 ± 52.3%). All had obtained a collegiate level of education (16.4± 3.2 years formal schooling) and were native speakers of American English. On average, the sample had 6.9 ± 6.6 years (range = 0 – 25 years) of formal, self-reported instrumental music training. All were paid for their time and gave informed consent in compliance with a protocol approved by the Institutional Review Board at Indiana University.

For each participant, we measured (i) vowel and CV phoneme categorization—to index gradient vs. categorical listening (Rizzi and Bidelman, 2024; Bidelman *et al*., 2025a), (ii) elliptical speech perception (discrimination, transcription) in clean and noise conditions (Quillet *et al*., 1998; Herman and Pisoni, 2003), and (iii) the QuickSIN (Killion *et al*., 2004), to assess SIN perception using a normed, external measure. Sounds were delivered binaurally through Sennheiser HD 280 circumaural headphones at a comfortable listening level (∼75 dB SPL) via custom scripts coded in MATLAB (v2024a; Natick, MA, USA).

### B. Phoneme categorization

We used vowel and consonant-vowel (CV) synthetic phoneme continua identical to our previous reports on speech categorization and SIN processing (e.g., Bidelman *et al*., 2019; Bidelman and Carter, 2023; Bidelman *et al*., 2025c). CVs contain stronger “intrinsic” features for identification (Pisoni, 1973; Xu *et al*., 2006) and are perceived more categorically than vowels (Pisoni, 1973; Altmann *et al*., 2014; Carter *et al*., 2022). Use of both continua allowed us to measure categorization for stimuli that differ in their salience of acoustic-phonetic mapping but also ensure categorization effects were not idiosyncratic to a specific class of phonemes.

#### 1. Vowel stimulus continuum

The vowel continuum was a synthetic 5-step vowel continuum spanning from “u” to “a” (for details see, Bidelman *et al*., 2013; Carter *et al*., 2022). Tokens were synthesized using a Klatt-based synthesizer coded in MATLAB (e.g., Klatt, 1980). Each token was separated by 5 equidistant steps acoustically based on first formant frequency (F1) (430 and 730 Hz). Individual vowel tokens were 100 ms in duration including 5 ms of ramping. Each token contained identical voice fundamental (F0), second (F2), and third formant (F3) frequencies (F0: 150, F2: 1090, and F3: 2350 Hz). Speech tokens were RMS amplitude normalized and ramped with cos^2^ windowing to reduce spectral splatter.

#### 2. Consonant-vowel stimulus continuum

The CV continuum consisted of a 5-step, stop-consonant /da/ to /ga/ sound gradient (e.g., Bidelman *et al*., 2019; Carter *et al*., 2022). CV tokens were adopted from natural production in Nath and Beauchamp (2012). Individual CV tokens were 350 ms in duration including 5 ms of ramping. Stimulus morphing was achieved by altering the F2 formant region in a stepwise fashion using the STRAIGHT toolbox (Kawahara *et al*., 2008).

#### 3. Categorization task

Categorization for both continua was measured using a visual analog scale (VAS) (Massaro and Cohen, 1983; Kapnoula *et al*., 2017; Rizzi and Bidelman, 2024; Bidelman *et al*., 2025a). The VAS paradigm required participants to mouse click a point along a continuous visual scale with endpoints labeled “u”/”da” and “a”/”ga” to report their percept. A VAS task was favored here because it allows for more continuous judgments in perception and distinguishes gradient vs. categorical listeners better than a conventional 2AFC approach (Massaro and Cohen, 1983; Kapnoula *et al*., 2017; Rizzi and Bidelman, 2024; Bidelman *et al*., 2025a). Use of the entire VAS scale was encouraged to avoid response bias. Participants were asked to respond as accurately and quickly as possible. There were 15 trials per token. Unless the participants had clarifying questions, no other instructions were provided regarding the use of the VAS scale (Kapnoula *et al*., 2017). Vowels and CVs were run in separate blocks (ordered randomly).

### C. Sentence materials

We measured perception of “elliptical speech” (Miller and Nicely, 1955) using the transcription and discrimination tasks described previously (Quillet *et al*., 1998; Herman and Pisoni, 2003). Raw stimulus materials consisted of 96 English sentences taken from the IEEE sentences (lists 1-10) (IEEE, 1969). Half of the utterances were spoken by a male speaker and the other half by a female speaker. Noise-masked versions of the sentences were created by mixing Gaussian noise with sentence materials at -5 dB SNR (Herman and Pisoni, 2003).

#### 1. Elliptical speech stimuli

Elliptical sentences were created through a manipulation of featural substitution (Miller and Nicely, 1955; Miller, 1956). For full details, see Herman & Pisoni (2003). Stops, fricatives, and nasal consonants in each of the keywords in each sentence were replaced with a different consonant that preserved the same manner and voicing features of the original consonant but transformed the place feature to an alveolar place of articulation. Liquids (/r/, /l/) and glides (/y/, /w/) were excluded. The “ellipsis” refers to the deletion of distinctive features and reduction of consonants to essentially five broader groupings (*p-t-k; f-*θ*-s-*L*; b-d-g; v-ð-z-*L*; m-n*) (Miller, 1956). Examples for an intact (original) and elliptical version of sentences are (1) below, with the keywords italicized:

a. A *wisp* of *cloud hung* in the *blue* air.—Intact A *wist* of *tloud hund* in the *dlue* air.—Elliptical
b. *Glue* the *sheet* to the *dark blue background*.— Intact *Dlue* the *seet* to eh *dart dlue datdround*.— Elliptical

#### 2. Discrimination (same/different) task

Sentence discrimination was measured for both elliptical and intact speech materials using a “same/different” task (Herman and Pisoni, 2003). Subjects were told they would be listening to pairs of English sentences containing familiar words. They were asked to label them as “same” if they perceived them as word-for-word and sound-for-sound identical or “different” if any of the words or speech sounds differed. There were 96 total sentence pairs evenly split between SNR conditions. Half of the sentences were elliptical speech and half were intact sentences containing the original place of articulation cues. All listeners received 8 types of sentence pair combinations denoted as *IiIj, EiEj, IiEj, EiIj, IiIi, EiEi, IiEi,* or *EiIi* (see Table 1 of Herman and Pisoni, 2003). In this nomenclature, intact and elliptical sentences are denoted as *I* and *E,* respectively, and sentence pairs with lexically identical words are marked with two lowercase *i’s*, whereas pairs of sentences that are lexically different are marked with a lowercase *i* and *j*. The critical conditions of interest were *IiEi* and *EiIi* sentences, which pair an intact and elliptical versions of an otherwise lexically identical sentence (only phonetic features are swapped). We expected these sentences would be perceived as sounding “different” in the clear but counterintuitively sound as the “same” in noise, if indeed listeners use broad phonetic categories for speech recognition under adverse listening conditions (Herman and Pisoni, 2003). *IiIj, EiEj, IiEj*, and *EiIj* sentences were expected to produce floor discrimination (i.e., always heard as “different”) given those pairings contained different word content—and thus elliptical processing would yield no phoneme confusions nor SNR benefit (Herman and Pisoni, 2003).

#### 3. Transcription task

We also measured participants’ ability to identify and transcribe elliptical sentences as a function SNR and place feature manipulations using an open-set transcription task (Quillet *et al*., 1998). There was a total of 192 sentences evenly divided by sentence type (intact vs. elliptical) and SNR (clean vs. noise) (i.e., 48 trials per condition). Subjects were instructed that they would hear a list of meaningful English sentences containing familiar words. For each of 3-5 keywords, a blank line was substituted in a text version of the sentence presented on screen. Listeners were instructed to type the words they heard as accurately as possible. Responses were logged via a MATLAB GUI interface.

### D. QuickSIN

The QuickSIN test (Killion *et al*., 2004) was used to obtained a normed assessment of perceptual SIN abilities. Participants heard six sentences embedded in four-talker noise babble, each containing five keywords. Sentences were presented at 70 dB HL. The signal-to-noise ratio (SNR) decreased parametrically in 5 dB steps from 25 dB SNR to 0 dB SNR. At each SNR, participants were instructed to repeat the sentence and correctly recalled keywords were logged. We computed their SNR loss by subtracting the number of recalled target words from 25.5 (i.e., SNR loss = 25.5 -Total Correct). The average from two lists was taken as the listeners’ QuickSIN score.

### E. Behavioral data analysis

#### 1. Categorization data

We estimated the slope of each listeners psychometric identification functions from their VAS labeling (Massaro and Cohen, 1983; Kapnoula *et al*., 2017; Rizzi and Bidelman, 2024). Identification scores (*P*) were fit with a sigmoid function (*P* = 1/[1+*e*^-β*1*(*x*-β*0*)^]) using Bayesian inference via the *psignifit* toolbox (Schütt *et al*., 2016). The relevant slope parameter (β*_1_*) reflects the steepness of the identification function where stronger, more discrete categorization produces larger β*_1_* slope values. Behavioral labeling speeds (i.e., reaction times [RTs]) were computed as the median response latency across trials for a given condition. RTs outside 250-2500 ms were deemed outliers (e.g., fast guesses, lapses of attention) and were excluded from the final analysis (Bidelman *et al*., 2013; Bidelman and Walker, 2017).

#### 2. Elliptical speech scoring

Sentence recognition in the transcription task was scored for keyword accuracy. In the case of elliptical speech sentences, scoring was determined based on whether the original word (i.e., prior to conversion to elliptical speech) was reported (Herman and Pisoni, 2003). Two independent raters manually scored the transcription data. Inter-rated agreement was excellent [*r* = 0.99, *p* < 0.0001]. Discrimination data were scored as the proportion of sentences reported as “same” for each of the 8 sentence pairs (Herman and Pisoni, 2003).

### F. Statistical analysis

Unless otherwise noted, we analyzed the dependent variables using linear mixed-effects models in R (version 4.2.2) (R-Core-Team, 2020) and the lme4 package (Bates *et al*., 2015) using maximum likelihood estimation. Sentence-level speech discrimination and transcription (%-same, %-keyword accuracy) were analyzed with fixed effects of SNR (clear, noise) and sentence type (intact, elliptical). Phoneme categorization measures (identification slope, RTs) were analyzed with fixed effects of continuum (vowels, CVs), and, in the case of RTs, token (Tk1-5). Subjects served as a random intercept. Discrimination and transcription scores were transformed via rationalized arcsine units (RAU) for the statistical models to account for ceiling/floor effects in percent correct data (Studebaker, 1985). Years of musical training served as a continuous covariate and was z-scored to position this variable on a similar scale as the other predictors^2^. Tukey-adjusted contrasts were used for multiple comparisons. Effect sizes are reported as partial eta squared (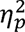) and degrees of freedom (*d.f.*) using Satterthwaite’s method. Spearman’s correlations assessed bivariate relationships between variables.

## III. RESULTS

### A. Phoneme categorization vs. QuickSIN

#### Phoneme identification

Phoneme categorization for CVs and vowels is shown in **Figure 1**. VAS responses showed large differences in listening strategy; some subjects produced highly binary responses and others provided more graded labelling across the continuum, consistent with previous reports (Rizzi and Bidelman, 2024; Bidelman *et al*., 2025a).

**Fig. 1.**
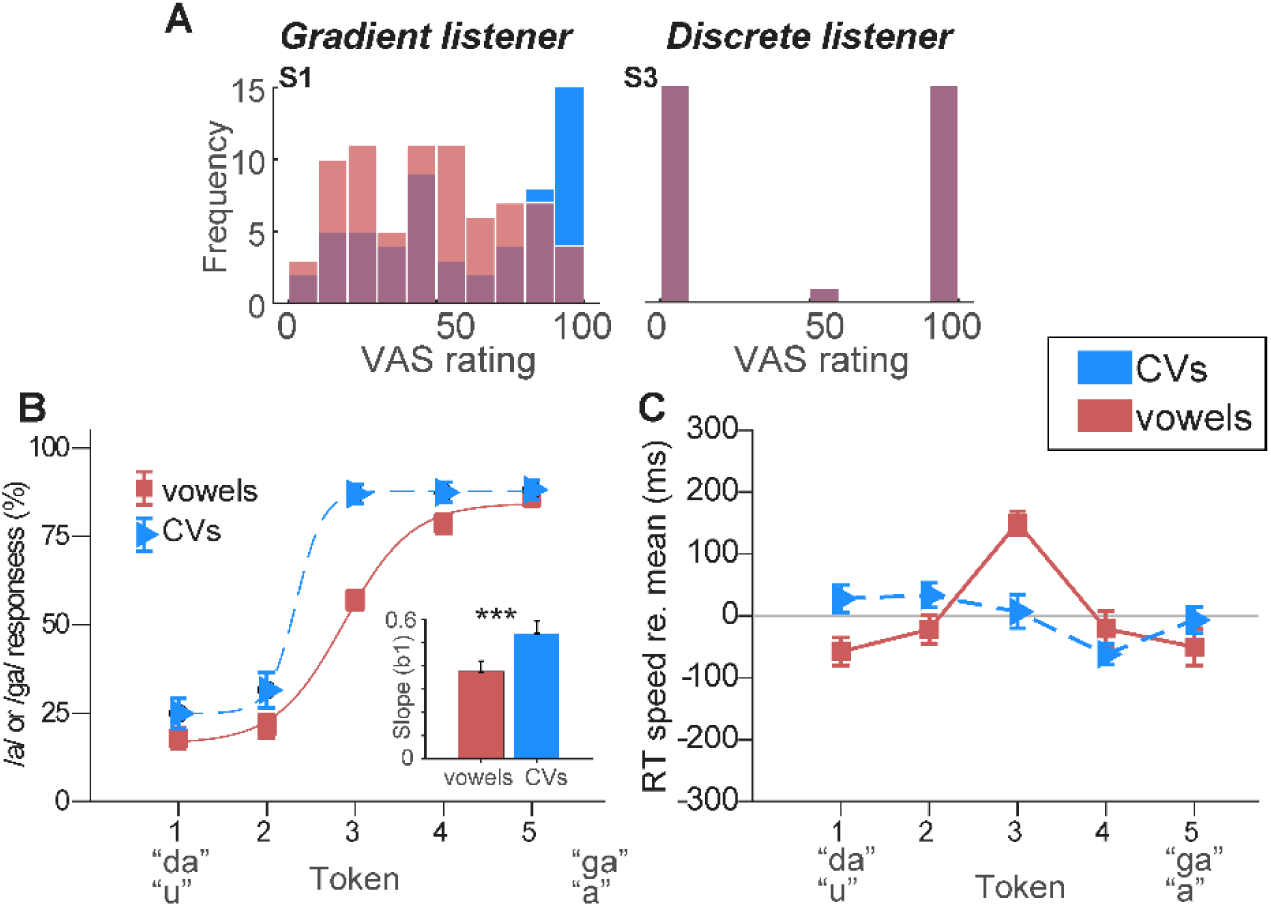
(Color online): Phoneme categorization. (**a**) Individual histograms of a representative gradient vs. discrete listener illustrating the distribution of VAS phonetic labeling for CV and vowel continua. Gradient listeners (e.g., S1) produce more continuous ratings across the continuum. Discrete listeners (e.g., S3) produce more binary categorization where responses lump near endpoint tokens. Categorization is highly stable within listener (similar labeling for CVs and vowels). (**b**) Phoneme identification curves derived from VAS labeling. Note the sharper, more discrete categorization for CVs compared to vowels. (**c**) Phoneme labeling speeds. RTs show the typical slowing near the perceptually ambiguous midpoint of the vowel (but not CV) continuum. RTs are plotted normalized to the global mean to highlight token-related changes across the two continua. errorbars = ± 1 s.e.m., \*\*\**p* < 0.0001.

**Fig. 1A** illustrates VAS responses from a representative “discrete” and “continuous” listener. Despite differences between individuals, responses were highly consistent for vowels and CVs *within*-listener (vowel and CV slope correlation: *r* = 0.73, *p* <0.0001). This suggests that the gradiency in listeners’ phoneme categorization was a stable perceptual trait (Rizzi and Bidelman, 2024; Bidelman *et al*., 2025a).

**Fig. 1B** shows psychometric identification functions derived from VAS labeling distributions. Identification slopes, reflecting the degree of categoricity in listeners’ response pattern, varied by phoneme type [*F_1,27_* = 19.92, *p* = 0.00012; 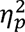 = 0.42] and were steeper for CVs compared to vowels, consistent with prior studies (Pisoni, 1973; Altmann *et al*., 2014; Carter *et al*., 2022; Bidelman *et al*., 2025a).

#### Labeling speeds

RT labeling speeds are shown in **Fig. 1C**. We normalized the RTs by subtracting the mean across tokens to highlight the relative changes in speed between continua and tokens (Bidelman *et al*., 2019). *A priori* contrasts revealed the hallmark slowing (i.e., inverted-V pattern) in labeling speeds near the ambiguous midpoint of the continuum for vowels [*t_240_* = 3.99, *p*=0.0001] (Pisoni and Tash, 1974; Bidelman and Walker, 2017; Carter and Bidelman, 2021). Thus, RTs were a positive function of stimulus uncertainty, increasing at the categorical boundary where labeling is least consistent (Pisoni and Tash, 1974). However, this slowing effect due to phonetic ambiguity was not observed for CVs [*t_240_* = 0.197, *p*=0.84], consistent with some prior speech categorization studies (Carter and Bidelman, 2021; Carter *et al*., 2022; Bidelman *et al*., 2025a). The differential pattern between vowel and CV categorization is well described by simple diffusion decision models that accumulate sensory evidence over time (Ratcliff and McKoon, 2008)^3^. CVs are heard more categorically than vowels which leads to more rapid evidence accumulation and informally faster RTs. Collectively, our phoneme categorization data support the notion that although CVs are generally perceived more categorically than vowels, certain listeners show inherently more gradient hearing of speech compared to others who show more categorical responses. Given the high consistency in vowel vs. CV labeling within subject, we pooled categorization measures for subsequent analyses to reduce the dimensionality of the data.

#### QuickSIN

We next assessed relations between phoneme categorization and QuickSIN performance. Several studies have shown links between the breadth of a listeners’ perceptual categories at the phoneme level and their SIN performance (e.g., Kapnoula *et al*., 2017; Myers *et al*., 2024; Rizzi and Bidelman, 2024; Bidelman *et al*., 2025a; Rizzi and Bidelman, 2025). We included musical training as a continuous covariate in these analyses given that (i) our sample included listeners with a wide spread of self-reported formal music training and (ii) prior studies suggest musicianship enhances both SIN processing in children and adults (Parbery-Clark *et al*., 2009b; Zendel and Alain, 2012; Anaya *et al*., 2016; Coffey *et al*., 2017; Mankel and Bidelman, 2018; Yoo and Bidelman, 2019; Bidelman and Yoo, 2020; Benítez-Barrera *et al*., 2022; Hennessy *et al*., 2022) and auditory category mapping (Elmer *et al*., 2012; Bidelman *et al*., 2014; Bidelman and Alain, 2015; Mankel and Bidelman, 2018; Bidelman and Walker, 2019; Ma *et al*., 2024). Indeed, QuickSIN scores were negatively correlated with self-reported musicianship in our sample [*r* = -0.44, *p*=0.0198] (**Fig. 2A**). Listeners with more extensive music engagement had better SIN performance, consistent with several previous studies (Hennessy *et al*., 2022; Maillard *et al*., 2023).

**Fig. 2.**
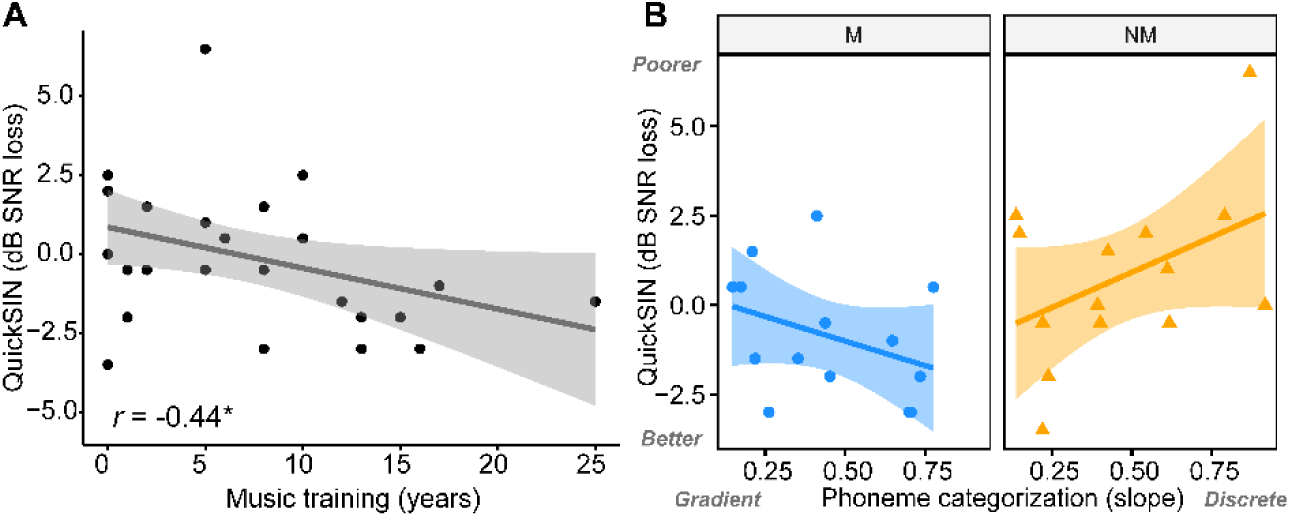
QuickSIN scores vary with musicianship and are related to phoneme categorization. (**A**) Listeners with more extended music training performed better on the QuickSIN. (**B**) Categorization slopes predict QuickSIN performance differentially with years of musical training. Better SIN is associated with steeper labeling slopes (i.e., more discrete phoneme perception) in listeners with more music (Ms), whereas in listeners with less music (NMs), better SIN relates to shallower slopes (i.e., more gradient/continuous phoneme perception). shading = 95% CI.

A multivariate linear model with QuickSIN as the dependent variable further revealed that SIN performance was related to phoneme categorization slopes but differentially depending on listeners’ music background and in opposite ways [slope * music interaction: *F_1,24_* = 4.43, *p* = 0.045; 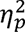] (**Fig. 2B**). Better QuickSIN performance was associated with more discrete phoneme labeling (i.e., steeper identification slopes) in musical listeners but more continuous perception (i.e., shallower slopes) in non-musical listeners. These data affirm the relation between phonetic perception and SIN perception (e.g., Myers *et al*., 2024; Rizzi and Bidelman, 2024; Bidelman *et al*., 2025a) but show this association varies with listeners’ music background.

### B. Elliptical speech perception

**Figure 3** shows mean discrimination and transcription results for clean and noise-degraded intact and elliptical speech sentences across listeners.

**Fig. 3.**
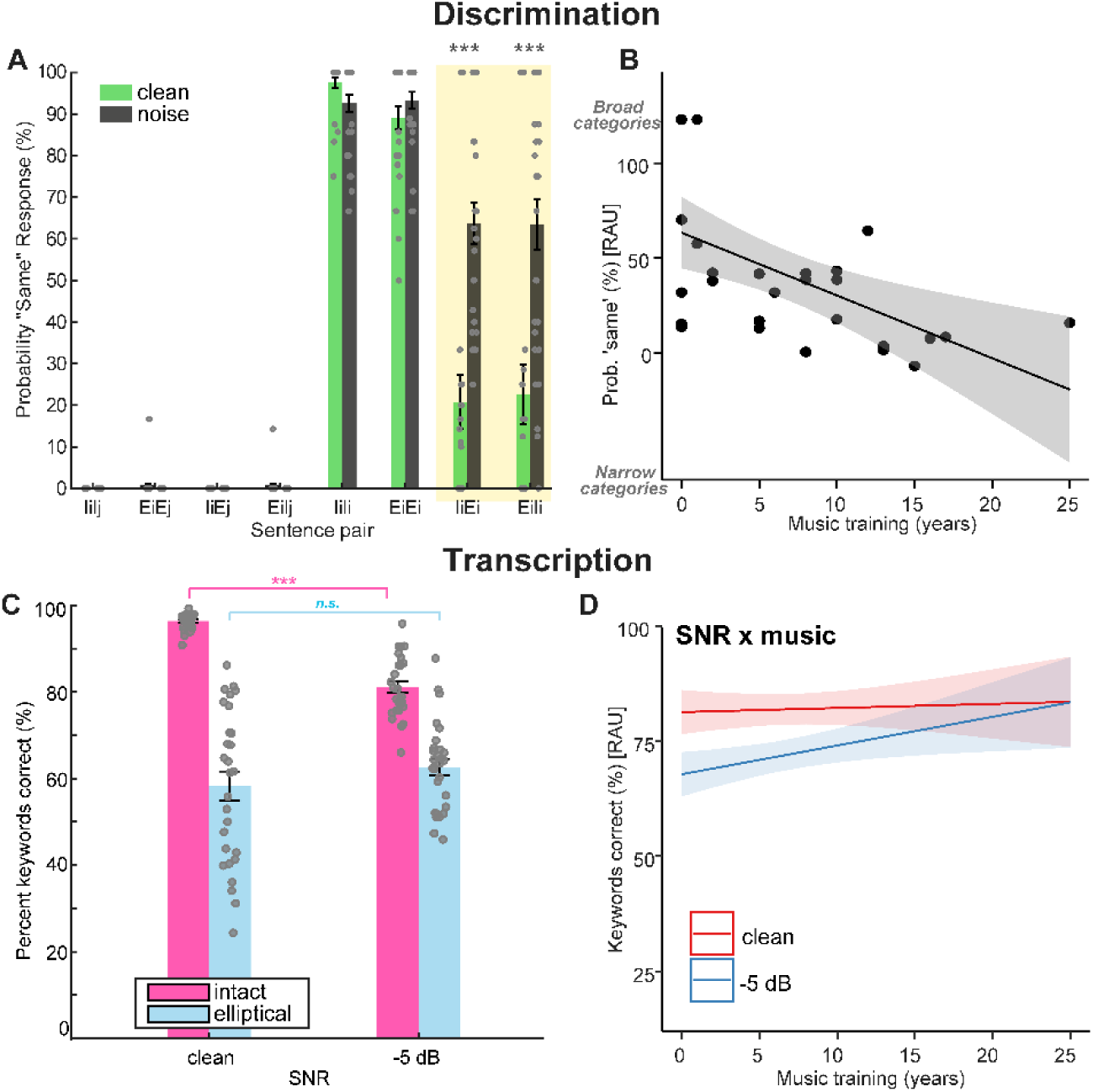
(Color online): Elliptical speech discrimination and transcription. (**A**) Discrimination for sentence pairs. Shading = critical test of elliptical speech benefit to SIN perception*. E* = elliptical sentences, *I*=intact sentences, *i/j* = unique lexical content. Ellipsis is undetectable under degraded listening conditions suggesting listeners use broad phonetic categories for SIN perception. (**B**) Main effect of music. Longer duration of music engagement was related to fewer “same” responses in sentence discrimination, i.e., less susceptibility to elliptical featural substitutions. (**C**) Sentence transcription results. Noise degrades speech recognition for intact but not elliptical sentences. (**D**) Marginal effects plot illustrating an SNR x music training interaction on transcription accuracy. Longer duration of music engagement was related to increased recognition for noise-degraded (but not clean) speech regardless of sentence type. Dots in panel A-C denote individual participant data. Shading = 95%CI. errorbars = ± 1 s.e.m., \*\**p* < 0.001, \*\*\**p* < 0.0001.

#### Sentence discrimination

As expected, sentences which differed in their lexical content (*IiIj, EiEj, IiEj, EiIj*) were always correctly perceived as “different” regardless of their elliptical quality **Fig. 3A**. These pairs were not included in the subsequent analyses given their trivially ceiling discrimination. A mixed-effect linear model conducted on discrimination scores of the remaining conditions revealed main effects of sentence pair [*F_3,20_* = 12.25, *p* <0.0001; 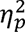 = 0.65], SNR [*F_1,20_* = 17.33, *p* = 0.00048; 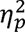 = 0.46], and years of music training [*F_1,20_* = 5.15, *p* =0.034; 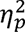 = 0.21]. There was also a critical sentence pair x SNR interaction [*F_3,20_* = 10.06, *p* =0.0003; 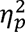 = 0.60]. Tukey-adjusted contrasts revealed the interaction was attributed to a differential effect of the ellipses on clean vs. noise-degraded speech. Physically identical sentence pairs (*IiIi, EiEi*) were correctly judged as being the same regardless of SNR (*ps* > 0.25). In contrast, the two critical pairs (*IiEi* and *EiIi;* **Fig. 3A**, shading) revealed that listeners perceived sentences with intact place of articulation features as being different than their elliptical counterparts when heard in the clear but more often as the same under noise degradation (*IiEi*: *p*=0.0001; *EiIi*: *p*<0.0001). This confirms the prediction that the ellipsis is less detectable under degraded listening conditions (Miller and Nicely, 1955; Quillet *et al*., 1998; Herman and Pisoni, 2003).

The main effect of music on sentence discrimination was a negative relationship. More extended music experience related to lower susceptibility to elliptical speech and therefore less frequent perceptual conflation of the transformed (i.e., swapped) phonemes when in noise (**Fig. 3B**). These findings again suggest “musicians” may have used narrower categories during sentence-level recognition whereas “nonmusicians” used broader phonetic categories (cf. Fig. 2).

#### Sentence transcription

Open-set transcription of elliptical sentences is shown in **Figure 3C**. A linear model revealed that while intact sentences expectedly yielded better performance than elliptical sentences overall [main effect of sentence type [*F_1,78_* = 258.62, *p* < 0.0001; 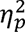 = 0.77], there was a critical sentence type x SNR interaction [*F_1,78_* = 30.88, *p* < 0.0001; 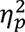 = 0.28]. Replicating Quillet et al. (1998), keyword identification for intact sentences declined markedly in noise (*p* < 0.0001). However, keyword identification for elliptical speech was remarkably resistant to noise degradation. In fact, recognition for elliptical speech tended to increase slightly (+4-5%) from clean to noise, though this effect did not reach significance (*p* = 0.08). Still, the usual maintenance of performance in noise suggests a perceptual benefit of ellipses on noise degraded speech perception (Quillet *et al*., 1998; Herman and Pisoni, 2003). Furthermore, we found a SNR x music interaction [*F_1,78_* = 3.82, *p* =0.051, 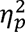 = 0.05] (**Fig. 3D**). Paralleling the QuickSIN results (see Fig. 2A), longer durations of music training were associated with better transcription performance for noise-degraded (but not clean) sentence materials across the board. These findings suggest: (i) elliptical transformations helped maintain intelligibility even as distortion of the signal increased and (ii) speech in noise performance of intact and elliptical speech varied with listener background.

### C. Relations between elliptical speech perception and QuickSIN

We next assessed relations between elliptical sentence and QuickSIN tasks under the premise that the breadth of a listeners’ perceptual categories might provide a common underlying construct to drive their SIN performance. Elliptical speech perception was also associated with QuickSIN performance but in opposite ways depending on listeners’ music experience. For elliptical speech discrimination, reporting “same” in the illusory elliptical conditions—indicative of broader category usage (i.e., yellow shading, Fig. 3)—related to *poorer* QuickSIN among “musicians” [*r_M_* = 0.71, *p* = 0.0047; *r_NM_* = -0.26, *p* = 0.36] (**Fig. 4A**). In contrast, better elliptical speech transcription scores—also indicative of broad category usage—related to *better* QuickSIN performance in “nonmusicians” [*r_M_* = 0.29, *p* = 0.31; *r_NM_*=-0.64, *p* = 0.014]. These findings suggest better SIN processing in musically-trained individuals related to narrower category use whereas improved SIN processing in musically-naïve individuals related to broader category use. These data are consistent with the notion that continuous modes of phonetic listening are more beneficial for SIN perception (cf. Rizzi and Bidelman, 2024; Bidelman *et al*., 2025a), especially in listeners with lower music expertise.

**Fig. 4.**
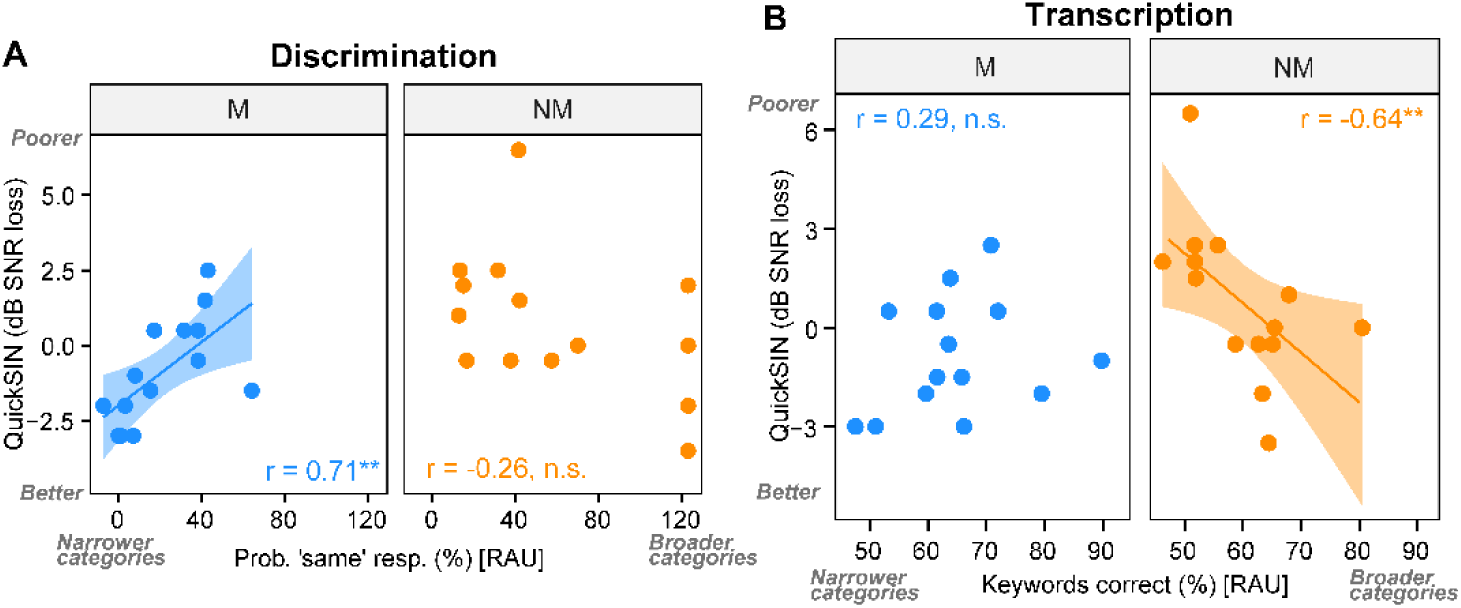
(Color online): Use of narrow/broad categories during sentence level perception is related to SIN perception but varies with musicianship. (**A**) Correlations between elliptical speech *discrimination* in noise (i.e., mean of critical IiEi/EiIi conditions; yellow shading, Fig. 3A) and QuickSIN. (**B**) Correlations between elliptical speech *transcription* in noise and QuickSIN. Musicians who use narrower categories at the sentence level show poorer SIN performance (i.e., higher QuickSIN scores), whereas nonmusicians who use broader categories show better SIN performance (i.e., lower QuickSIN scores). Shading = 95% CI. \*\**p* < 0.01.

## IV. DISCUSSION

The present study assessed relationships between auditory categorization for speech sounds (CV, vowels) and speech-in-noise recognition to test the premise that gradiency/categoricity at the phoneme and/or sentence level might account for noise-degraded speech listening skills. To test this hypothesis, “elliptical speech” (Miller and Nicely, 1955; Miller, 1956) was used to assess listeners’ propensity for perceiving featural substitutions within sentences that render them meaningless under clear conditions but surprisingly improves their recognition under noise (Quillet *et al*., 1998; Herman and Pisoni, 2003). Our findings show that listeners who perceived speech using more gradient phonetic categories at the token level and broader equivalency classes at the sentence level enjoy larger benefit in perceiving SIN. However, this effect is highly dependent on listeners’ music background.

Using standard phoneme categorization tasks, we confirmed substantial variation in how listeners labelled isolated speech sound continua. Replicating prior work, CVs were perceived more categorically than vowels (Pisoni, 1973; Altmann *et al*., 2014; Carter *et al*., 2022; Bidelman *et al*., 2025a), consistent with the notion that stops carry stronger intrinsic acoustic features upon which to make category judgments (Xu *et al*., 2006; Bidelman *et al*., 2020). More importantly, listeners varied in their degree of gradiency in phoneme labeling; some listeners showed VAS response patterns representative of “categorical” listening strategies while others showed “continuous” perception. Despite such differences between individuals, responses were highly consistent for vowel and CV tokens *within*-listener. This suggests that being a discrete vs. continuous listener is a stable perceptual trait that does not depend on the speech materials nor the specific acoustic-phonetic continuum (Rizzi and Bidelman, 2024; Bidelman *et al*., 2025a). Some listeners appear to be more gradient/continuous in hearing out phonetic features compared to others who show a more discrete/categorical mode of listening (Kapnoula *et al*., 2017; Apfelbaum *et al*., 2022; Rizzi and Bidelman, 2024; Bidelman *et al*., 2025a).

### A. Elliptical speech reveals SIN processing depends on how listeners deploy category-relevant information

Results from our speech recognition tasks closely replicate prior studies on elliptical speech perception (Miller and Nicely, 1955; Quillet *et al*., 1998; Herman and Pisoni, 2003). While recognition of clear *intact* speech expectedly declined in noise, discrimination performance on *elliptical* speech remained largely invariant to noise degradation. Sentences with intact place of articulation and their elliptical version were labeled *different* when heard in the clear but were labeled as the *same* when heard under noise (cf. Herman and Pisoni, 2003). This indicates that listeners failed to detect the featural substitutions and thus heard elliptical and intact speech as sounding identical under noise. Similar results were observed for sentence transcription. While elliptical speech transcription was poorer than intact transcription overall, keyword identification for elliptical speech remained robust to noise, and even increased slightly at the lower SNR. Both sentence discrimination and transcription data confirm that despite their featural swaps, elliptical speech remained surprisingly invariant to noise and was heard as sounding more like clear speech under conditions of acoustic degradation.

One interpretation of these findings is that when the acoustic-phonetic input is impoverished, listeners could merge certain phonetic contrasts into broader equivalence classes to recognize speech (Quillet *et al*., 1998). Thus, even if a listener fails to perceive the correct place of articulation, they might still be able to recognize the intended words correctly by using broader manner equivalence classes (Shipman and Zue, 1982). This form of perceptual reorganization that exploits broader category use may provide reliable support for word recognition and lexical access under degraded listening conditions where only partial acoustic-phonetic information is available (Quillet *et al*., 1998; Herman and Pisoni, 2003). While the present results were for normal-hearing listeners, even cochlear implant users—whose coarse acoustic input provides sparse sensory representations—benefit from using broader perceptual categories in noise as measured in elliptical speech experiments (Herman and Pisoni, 2003).

Broader category use with different forms of signal degradation could also reflect increased reliance on top-down, “predictive coding” (Friston, 2010) in order to guide speech perception in the face of impoverished sensory representations at the input (Herman and Pisoni, 2003; Sohoglu *et al*., 2012; Cope *et al*., 2017). In this regard, our data are reminiscent of the phenomenon of auditory induction and phonemic restoration when listeners still correctly perceive target speech even when it is momentarily interrupted by foreground noise (Warren, 1970; Warren *et al*., 1972). Perceptual continuity is also maintained even when the signal is physically absent (i.e., silent) during noise (Warren, 1970; Riecke *et al*., 2007; Riecke *et al*., 2009; Bidelman and Patro, 2016). This suggests the auditory system can fill in and perceptually restore missing segments occluded from the auditory scene through “top-down” inference. Under this interpretation, a broad category perceiver might be more liable to collapse acoustic-phonetic variation and thus more tolerant of distortions as demonstrated by our elliptical speech tasks.

Of particular interest were relations between listeners’ deployment of perceptual categories at phoneme and sentence levels and their SIN processing. Several reports have implied links between isolated phoneme categorization and successful SIN recognition, though the direction of such effects has been equivocal (Gifford *et al*., 2014; Kapnoula *et al*., 2017; Bidelman *et al*., 2020; Bidelman and Carter, 2023; Myers *et al*., 2024; Rizzi and Bidelman, 2024; Bidelman *et al*., 2025a; Rizzi and Bidelman, 2025). Gradient listening strategies (i.e., employing broader category usage) might confer a perceptual advantage if listeners are able to encode and maintain within-category acoustic details and thus better “hedge” their bets under degraded speech conditions (Toscano *et al*., 2010; Kapnoula *et al*., 2017; Apfelbaum *et al*., 2022; Rizzi and Bidelman, 2024). Alternatively, discrete/categorical listening could make a listener more decisive in perceiving speech (Bidelman *et al*., 2020; Bidelman and Carter, 2023) by effectively downsampling the acoustic space to retain only high-level phonetic category information that is more resistant to noise interference (Helie, 2017; Bidelman *et al*., 2020). Our correlations between both phoneme labeling and QuickSIN (Fig. 2) as well as elliptical speech perception and QuickSIN (Fig. 4) provide converging evidence that affirms the connection between auditory category usage and SIN processing. However, we find the direction in how category usage maps to improved SIN perception varies critically with listeners’ musical background.

### B. Auditory expertise modulates category usage and its link to SIN processing

Across our sample, listeners’ years of musical training predicted elliptical speech performance and its related noise benefit. That “nonmusicians” showed more susceptibility to elliptical speech than “musicians” (Fig. 3B) suggests that individuals with less auditory expertise might employ broader phonetic categories (or heavier top-down inference) during speech perception. This would allow them to obtain greater benefit from phoneme swaps during elliptical sentence manipulations. In contrast, listeners with more extensive music engagement seem to show less susceptibility to elliptical speech, suggesting they use narrower categories in forming their perceptual decisions and coping with signal uncertainty.

With regard to SIN processing, several studies suggest musicians might have a perceptual benefit in “cocktail party” listening scenarios (for reviews see Alain *et al*., 2014; Coffey *et al*., 2017; Hennessy *et al*., 2022; Bidelman *et al*., 2025b). Across the lifespan, self-reported musicians outperform nonmusicians in perceiving degraded speech under noise and reverberation (Parbery-Clark *et al*., 2009a; Bidelman and Krishnan, 2010; Parbery-Clark *et al*., 2011; Zendel and Alain, 2012; Kraus *et al*., 2014; Swaminathan *et al*., 2015; Anaya *et al*., 2016; Clayton *et al*., 2016; Brown *et al*., 2017; Deroche *et al*., 2017; Du and Zatorre, 2017; Mankel and Bidelman, 2018; Torppa *et al*., 2018; Yoo and Bidelman, 2019; Bidelman and Yoo, 2020; Lo *et al*., 2020; Brown and Bidelman, 2022) (but see Madsen *et al*., 2019; Whiteford *et al*., 2025). Considering music as a continuous variable, we found that listeners with more extended self-reported music training performed better on the QuickSIN (Fig. 2) consistent with several previous studies (Parbery-Clark *et al*., 2009b; Zendel and Alain, 2012; Fuller *et al*., 2014; Başkent *et al*., 2018; Mankel and Bidelman, 2018; Yoo and Bidelman, 2019; Hennessy *et al*., 2022; Maillard *et al*., 2023).

Music-related differences were also apparent in the direction by which category usage at the phoneme (Fig. 2) and sentence (Fig. 4) levels were related to SIN. For example, “musicians” who utilized broader categories during elliptical speech showed *poorer* QuickSIN performance. In contrast, broader category usage in “nonmusicians” was associated with *better* QuickSIN performance. Thus, while gradiency might benefit hearing in noise for musically lay listeners (present study; Rizzi and Bidelman, 2024; Bidelman *et al*., 2025a), SIN processing in highly trained ears might instead benefit from discrete hearing strategies that employ narrower category use.

### C. Mechanisms for differential categorization-SIN relations among musicians and nonmusicians

What might be the basis for the differential effects of music demographics? Several pieces of evidence suggest that musicians may fundamentally approach auditory perceptual tasks differently than nonmusicians, including categorization and SIN. Musicians show enhanced classification of speech and musical sounds as indicated by steeper and faster speech identification (Elmer *et al*., 2012; Zuk *et al*., 2013; Bidelman *et al*., 2014; Bidelman and Alain, 2015; Mankel and Bidelman, 2018; Bidelman and Walker, 2019; Mankel *et al*., 2020; Ma *et al*., 2024). The present results using elliptical speech generally agree with these earlier findings from isolated phoneme identification experiments and reveal musicians may also use narrower, more discrete categories at the sentence level.

Our data are broadly consistent with the *distinctness hypothesis* which states that differences in the robustness and clarity of representation can cause success or failure in readout of high-order linguistic information (Elbro, 1996). Lexical quality also has direct consequence for more complex, downstream processes like comprehension (Perfetti, 2007). Thus, listeners with higher quality phonetic representations might perform better in SIN comprehension even when the quality of those representations is compromised by noise. Musicians, by virtue of their more robust auditory neural encoding of speech (e.g., Baumann *et al*., 2008; Bidelman *et al*., 2014; Bidelman and Alain, 2015; Bidelman and Walker, 2019; Kim *et al*., 2025), may enjoy higher quality and more “distinct” phonetic representations. Indeed, EEG studies have shown that musicians have larger Euclidean distance between their neural responses to different musical chords, leading to higher information transfer and reduced behavioral confusions of sound categories, indicative of narrower category use (Bidelman *et al*., 2011; MacLean *et al*., 2024). The same might apply to speech perception (e.g., Elmer *et al*., 2014; Varnet *et al*., 2015; Bidelman and Walker, 2019). Words that are distinct are more easily remembered and retrieved from memory which may facilitate better abstraction during higher level linguistic processing (Elbro *et al*., 1994). At the opposite extreme, more *in*distinct speech representations can lead to dyslexia and reading difficulties (Elbro *et al*., 1994; Elbro, 1996) which are often comorbid with SIN deficits (Nittrouer *et al*., 2018; Van Hirtum *et al*., 2021). Elbro (1996) further describes a system with more “distinctness” as being more robust and insensitive to disturbances like noise (Elbro, 1996, p.468). Thus, poorer, more incomplete, and underspecified representations of the speech signal may lead to lower distinctness at the phoneme level and poorer sentential QuickSIN performance as observed here in our inter-task correlations.

“Musicians” and “nonmusicians” also differ in their neural processing of speech and where categories are decoded in the brain. Whereas musicians show stronger category representation in early auditory cortical areas, nonmusicians relegate category processing to higher-order, inferior-frontal regions including Broca’s area (Bidelman and Walker, 2019). Neural coupling within the fronto-temporal pathways is critical for both successful speech categorization (Binder *et al*., 2004; Myers *et al*., 2009; Du *et al*., 2014; Alho *et al*., 2016; Bidelman and Walker, 2019; Carter and Bidelman, 2021) and degraded speech processing (Eisner *et al*., 2010; Bidelman and Dexter, 2015; Bidelman *et al*., 2018; Carter and Bidelman, 2021; Schelinski and von Kriegstein, 2023). Thus, the observed group differences may result from heavier reliance on top-down predictive coding vs. stronger bottom-up driven signal processing to extract stimulus-specific features of the speech signal (Herman and Pisoni, 2003; Sohoglu *et al*., 2012; Cope *et al*., 2017; Morillon and Baillet, 2017). Several studies have shown that musicianship indeed alters the balance between bottom-up vs. top-down processing during speech perception and lexical semantic tasks (Loui, 2016; Gagnepain *et al*., 2017; Bidelman and Walker, 2019; MacLean *et al*., 2025; Manting *et al*., 2025). In a sense, narrower phonetic category usage and enhanced SIN perception in musical listeners might reflect more “bottom-up” dominant processing given their higher quality of neural representation across multiple levels of the auditory system hierarchy (Weiss and Bidelman, 2015; MacLean *et al*., 2024).

## V. CONCLUSIONS

Elliptical speech stimuli offer a novel window into potential links between auditory categorization and noise-degraded speech perception (cf. Kapnoula *et al*., 2017; Bidelman *et al*., 2020; Kapnoula *et al*., 2021a; Bidelman and Carter, 2023; Myers *et al*., 2024; Rizzi and Bidelman, 2024; Bidelman *et al*., 2025a; Rizzi and Bidelman, 2025). Overall, our findings suggest: (i) phoneme categorization and SIN skills are related; (ii) the use of broad acoustic-phonetic categories can positively benefit noise-degraded speech perception (Quillet *et al*., 1998; Herman and Pisoni, 2003); (iii) successful SIN perception varies with how a listener forms perceptual equivalences in speech and this depends on their music background.

Different mechanisms might support how listeners approach noise-degraded listening scenarios dependent on their auditory expertise. Those with less music training could use broader phonetic categories to achieve robust SIN recognition, which might reflect increased reliance on “top-down” inference to maintain more flexible perception amidst noise (Herman and Pisoni, 2003; Sohoglu *et al*., 2012; Cope *et al*., 2017). In contrast, listeners with extensive music training may use narrower phonetic categories to achieve robust SIN recognition, which could reflect more “distinctness” in sensory representations that are insensitive to disturbances like noise (Elbro, 1996; Bidelman *et al*., 2020).

## ACKNOWLEDGMENTS

Work supported by the NIH/NIDCD R01DC016267 (awarded to G. M. B.). The authors thank Alexandra Doty for assistance in recruitment and data collection.

## AUTHOR DECLARATIONS

### Conflict of Interest

The authors have no conflicts of interest to disclose.

### Ethics Approval

Participants gave written, informed consent in compliance with a protocol approved by the Institutional Review Board at Indiana University (#14860).

### Data availability statement

The data that support the findings of this study are available from the corresponding author upon reasonable request. Modelling code is available on GitHub: https://github.com/bidelmanLab/diffusion-model-Pisoni-Tash-1974

## Footnotes

1 We note musician SIN benefits are not observed consistently in all studies (see Madsen *et al*., 2019; Escobar *et al*., 2020).

2 For the purpose of visualizing interactions with continuous variables, the data are often plotted with participants split into two groups based on a median split of music training (sample median = 5.5 years): “musicians” (M; n=14) and “nonmusicians” (NM; n=14) (Mankel and Bidelman, 2018; MacLean *et al*., 2024) (for similar definitions, see Zhang *et al*., 2020)

3 Pisoni & Tash (1974) reported slower RTs near the midpoint of their CV /ba/-/pa/ continuum (their Fig. 3). An inverted-V pattern in RTs during phoneme labeling is expected when tokens have increased stimulus ambiguity relative to adjacent steps along the continuum. Indeed, identification at the middle of Pisoni & Tash’s stimulus set showed non-ceiling labeling (∼75%). This suggests their listeners heard the midpoint token with some ambiguity, leading to the prolonged RT near the category boundary (see their Fig. 3). This mirrors the invert-V pattern we find for vowels, which elicit weaker, more ambiguous categorical percepts and similarly show a slowing of the RT near midpoint. When tokens are heard with an unambiguous category membership and labelled without confusion, RTs should be equally fast. Indeed, the flat RT pattern we find for our /da/-/ga/ continuum (e.g., “CVs”, Fig. 1) suggests each token was heard in a highly categorical manner with little ambiguity. Note too the more step like, floor/ceiling %-responses for CVs which confirms they were heard more ambiguously. In other words, the psychometric function must show performance approaching ∼50% at its inflection point to show a V-shaped slowing in RTs. Both labeling and RT differences for CVs vs. vowels are easily explained by simple decision models that assumes a noisy decision process which accumulates evidence over time (Ratcliff and McKoon, 2008). See model code and simulations on GitHub (https://github.com/bidelmanLab/diffusion-model-Pisoni-Tash-1974).

## Notes

### Competing Interest Statement

The authors have declared no competing interest.

### Summary of Updates

New analysis and results. Revised figures.

